# BenchDrop-seq: a microfluidics-free platform for benchtop single-cell long-read RNA sequencing

**DOI:** 10.64898/2026.03.12.706999

**Authors:** Jamie Bregman, Calen Nichols, Rajeev Ramisetti, Avi Srivastava

## Abstract

Single-cell long-read RNA sequencing enables direct measurement of full-length transcripts but has remained difficult to deploy at scale due to reliance on microfluidic barcoding, specialized instrumentation, and high per-cell cost. Here we present BenchDrop-seq, a benchtop platform for single-cell long-read transcriptomics that leverages particle-templated partitioning for single-cell molecular barcoding and couples this workflow to Oxford Nanopore sequencing for full-length transcript capture. By integrating established bead-based partitioning chemistry with long-read sequencing and a dedicated open-source analysis pipeline for barcode recovery, alignment, and transcript quantification, BenchDrop-seq enables isoform-resolved measurements from thousands of individual cells using standard laboratory equipment. We validate the platform in both a homogeneous cell line and a heterogeneous primary tissue, demonstrating high barcode recovery, accurate gene-level quantification, and reproducible detection of cell-type-specific transcript usage that is not readily accessible to short-read assays. Together, BenchDrop-seq establishes a practical and accessible framework for single-cell long-read RNA sequencing, lowering experimental barriers while enabling transcript-level analyses in routine single-cell experiments.

## Introduction

Alternative splicing expands the functional output of the genome by enabling individual genes to generate multiple transcript isoforms with distinct structural and regulatory properties^1^. Differences in isoform expression contribute to cellular identity and function^2^, while aberrant splicing, including shifts in isoform usage within specific cellular subsets, has been associated with diverse disease contexts^3^. Accurate measurement of transcript structure and usage at single-cell resolution, therefore, remains a central technical challenge for dissecting post-transcriptional regulation and cellular heterogeneity.

Bulk RNA sequencing averages transcriptomic signals across large cell populations, masking cell-to-cell variability in transcript structure and usage^4^. Single-cell RNA sequencing (scRNA-seq) overcomes this limitation by profiling individual cells, but most widely adopted scRNA-seq assays rely on short-read sequencing and capture only the 3’ or 5’ ends of transcripts^5,6^. This end-biased sampling collapses transcript structure and obscures isoform-specific information, forcing transcript diversity to be aggregated into gene-level counts^7^. As a result, differences in transcript usage across cells must be inferred indirectly or are lost altogether, particularly for genes with multiple isoforms, overlapping genomic loci, or long transcriptional units^8^. These constraints fundamentally limit short-read scRNA-seq’s ability to resolve gene and transcript-level heterogeneity within individual cells.

Long-read sequencing technologies address these limitations by generating reads that span entire transcript molecules, enabling direct observation of transcript structure without computational reconstruction^9^. When combined with single-cell barcoding, long-read sequencing enables, in principle, isoform-resolved single-cell transcriptomics. In practice, however, existing single-cell long-read workflows remain difficult to deploy. Most approaches rely on droplet-based microfluidic barcoding systems that require specialized instrumentation, incur high per-cell costs, and limit scalability^10–12^. Alternatively, combinatorial indexing strategies can increase throughput but often introduce experimental complexity or barcode ambiguity, compromising full-length transcript recovery^13^. In addition, long-read platforms yield fewer reads per sequencing run than short-read technologies^14^, further increasing cost and constraining experimental design. Together, these practical barriers have limited the widespread adoption of single-cell long-read transcriptomics despite its conceptual advantages.

To make isoform-resolved single-cell transcriptomics practical in routine laboratory settings, we developed BenchDrop-seq, an integrated experimental and computational platform for single-cell long-read RNA sequencing that operates entirely on a benchtop using standard laboratory equipment. BenchDrop-seq replaces microfluidic-based single-cell barcoding with particle-templated instant partitioning^15^ to achieve rapid single-cell partitioning and molecular barcoding, and couples this workflow to Oxford Nanopore sequencing for full-length transcript capture. The experimental protocol is paired with Bagpiper, an open-source analysis pipeline for barcode recovery, read alignment, and quantitative gene- and transcript-level analysis of single-cell long-read data. Together, these components enable recovery and quantification of full-length transcripts from thousands of individual cells without specialized instrumentation or proprietary software.

## Results

### Design and implementation of the BenchDrop-seq platform

We developed BenchDrop-seq as an integrated experimental and computational platform for single-cell long-read RNA sequencing that operates entirely on the benchtop without reliance on microfluidic instrumentation (Fig. 1; Material & Methods). The platform combines particle-templated instant partitioning for single-cell barcoding with Oxford Nanopore sequencing to enable recovery and quantification of barcoded full-length transcripts from thousands of individual cells.

**Figure 1.**
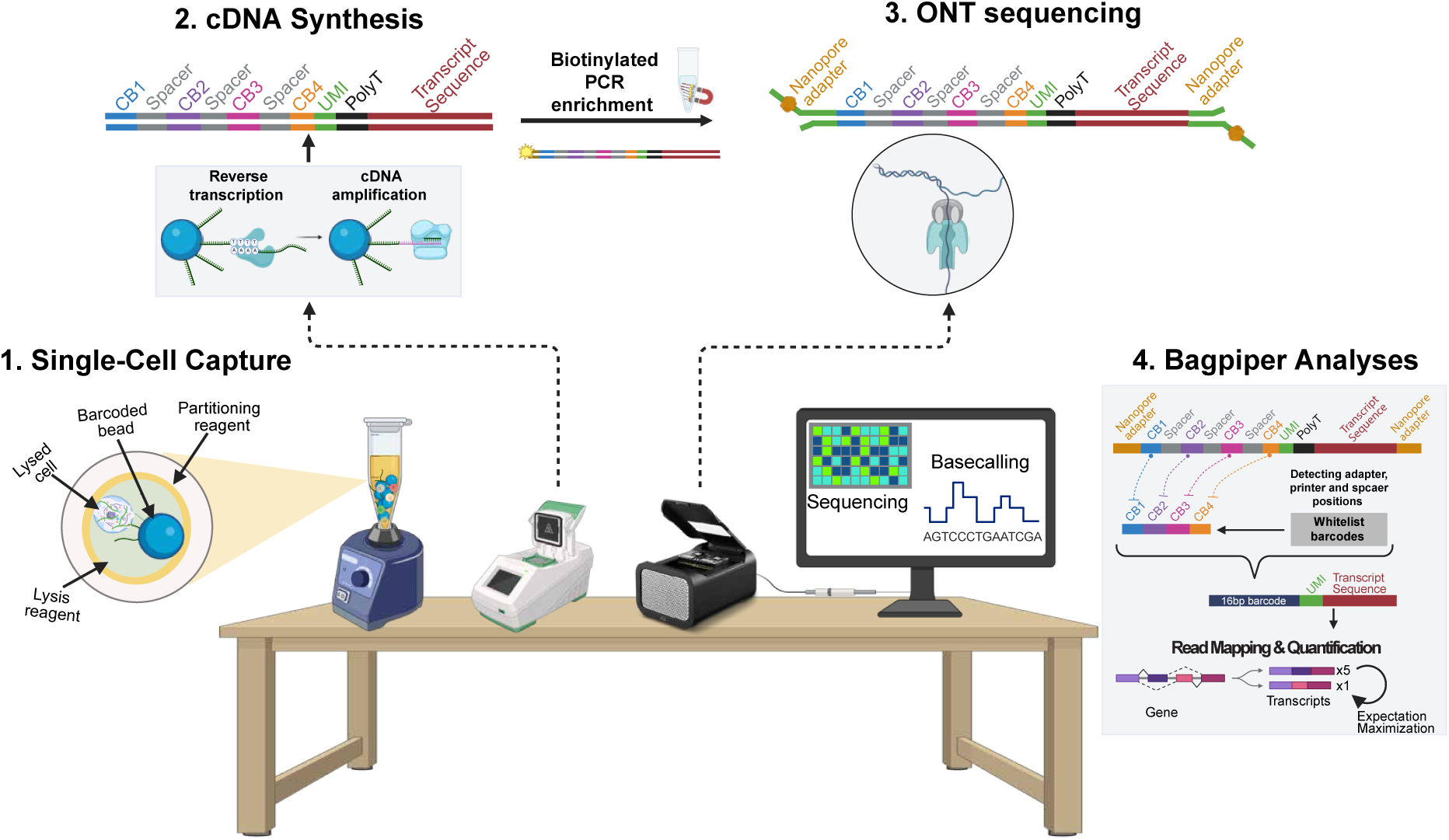
Overview of the BenchDrop-seq experimental workflow and Bagpiper analysis pipeline. BenchDrop-seq combines particle-templated single-cell barcoding with long-read sequencing and an integrated computational framework to enable benchtop single-cell long-read RNA sequencing and transcript quantification. (1) Single-cell capture and barcoding. Individual cells are lysed and partitioned into emulsions containing barcoded polyacrylamide beads using particle-templated instant single-cell partitioning. Released polyadenylated RNA molecules hybridize to bead-bound oligonucleotides carrying concatenated cellular barcodes and unique molecular identifiers. (2) cDNA synthesis and amplification. Reverse transcription and whole-transcriptome amplification are performed on bead-bound RNA, generating barcoded full-length cDNA molecules. Biotinylated PCR enrichment is used to selectively recover amplified cDNA for downstream library preparation. (3) Long-read sequencing. Amplified single-cell cDNA is prepared using Oxford Nanopore ligation-based chemistry and sequenced to generate long reads spanning cellular barcodes, unique molecular identifiers, and transcript sequences. Basecalling converts raw signal data into nucleotide sequences. (4) Bagpiper analyses. The Bagpiper pipeline identifies barcode and spacer sequences within long reads, assigns reads to whitelisted cellular barcodes, and aligns reads to the transcriptome. Gene- and transcript-level abundances are estimated using isoform-aware probabilistic assignment and expectation-maximization-based quantification.

In the BenchDrop-seq experimental workflow, cells are rapidly partitioned into emulsions containing barcoded polyacrylamide beads using vortex-based mixing. Following cell lysis, polyadenylated RNA molecules hybridize to bead-bound oligo(dT) carrying cellular barcodes and unique molecular identifiers (UMIs), enabling reverse transcription and molecular indexing at single-cell resolution. Full-length cDNA is subsequently amplified and prepared for Nanopore sequencing using ligation-based chemistry, generating long reads that span complete transcripts (Supplementary Fig. 1A; Material & Methods). This workflow replaces droplet-based microfluidic partitioning with a simple benchtop procedure while preserving single-cell resolution and full-length transcript coverage.

BenchDrop-seq was designed in parallel with a dedicated computational analysis layer to ensure reproducible processing of single-cell long-read sequencing data. As a core component of the platform, we developed Bagpiper, an open-source pipeline for barcode recovery, read alignment, and quantitative gene- and transcript-level analysis. Bagpiper identifies cellular barcodes directly from long reads using adaptive local alignment, aligns reads to reference transcriptomes using minimap2^16^, and performs isoform-aware gene and transcript quantification using an expectation-maximization framework tailored to the error characteristics of long-read sequencing in minutes (Supplementary Fig. 1B-C; Material & Methods).

Together, the experimental workflow and Bagpiper analysis pipeline constitute an end-to-end platform for single-cell long-read transcriptomics that relies on neither microfluidic barcoding systems nor proprietary software. While several approaches exist for generating^10,17–19^ and analyzing^20–24^ long-read transcriptomic data, many are optimized for bulk inputs or require specialized library structures and instrumentation. In contrast, BenchDrop-seq emphasizes simplicity, accessibility, and integration, providing a unified platform for microfluidics-free single-cell long-read RNA sequencing.

In the following sections, we benchmark BenchDrop-seq against matched short-read single-cell and bulk RNA-seq datasets to assess alignment performance, quantitative accuracy, and transcript-level resolution. Practical considerations, including protocol duration, reagent cost, scalability across single-cell RNA-sequencing workflows, and computational resource requirements, are detailed in the Materials and Methods & Supplementary Fig. 1, Table 1.

### Benchmarking gene-level quantification in a controlled cell system

We first evaluated BenchDrop-seq using human K562 erythroleukemia cells, a homogeneous system widely used to benchmark single-cell transcriptomic assays. In this system, we focused on benchmarking gene-level quantification accuracy using pseudobulk aggregation, enabling comparison to bulk RNA-seq measurements. We processed 2,468 high-quality cells and generated BenchDrop-seq single-cell long-read libraries and 3′-biased short-read libraries from the same barcoded cDNA pool (Fig. 2A-K562; Material & Methods), enabling direct platform-level comparison under controlled experimental conditions.

**Figure 2.**
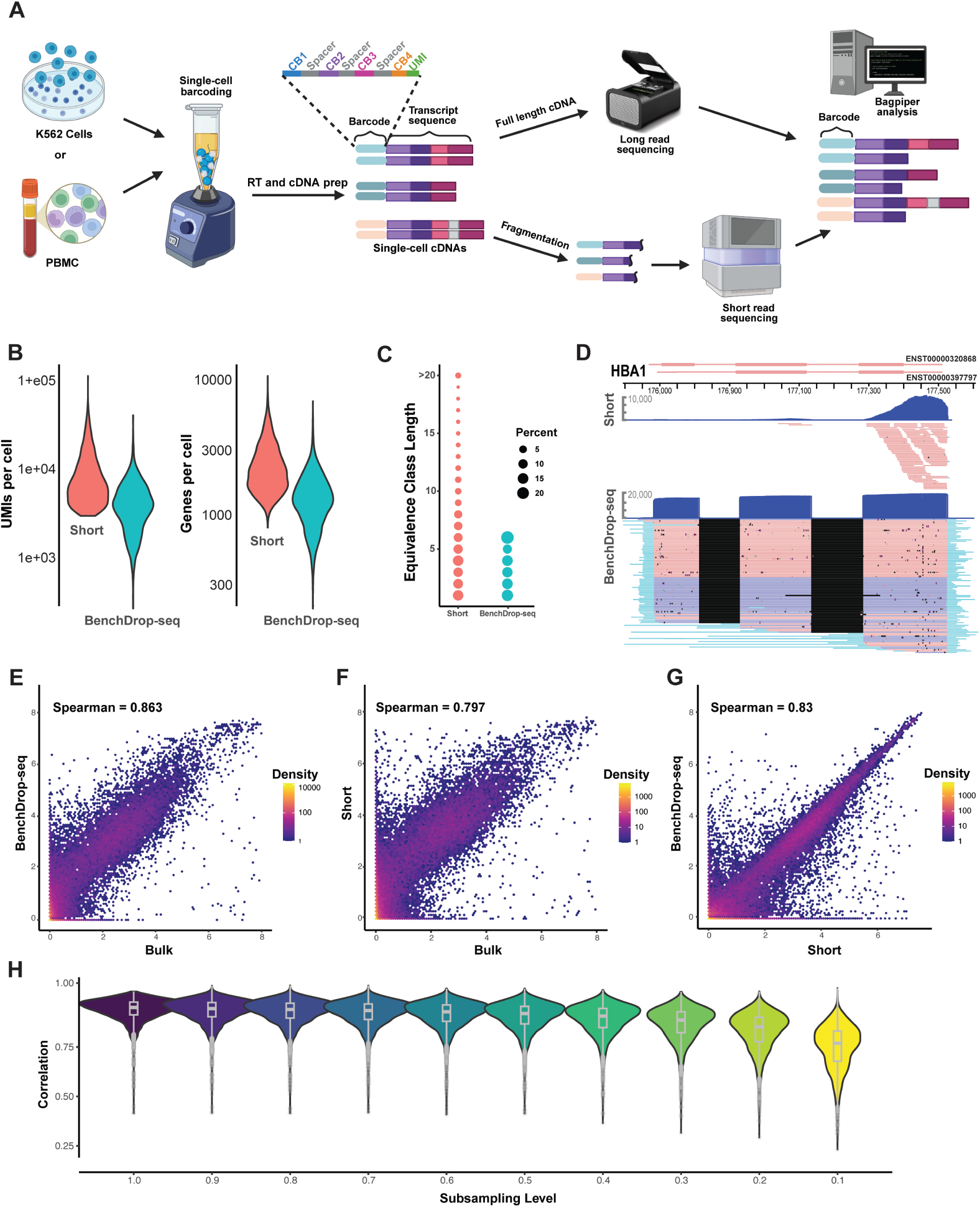
BenchDrop-seq enables accurate gene-level quantification from single-cell long-read data in a controlled cell system. Benchmarking in K562 cells demonstrates that single-cell long-read sequencing with BenchDrop-seq reduces transcript-assignment ambiguity and achieves robust gene-level concordance with bulk RNA-seq at modest sequencing depth. (A) Schematic overview of the BenchDrop-seq experimental workflow and analysis strategy. K562 cells or PBMCs are barcoded using particle-templated instant partitioning, followed by reverse transcription and cDNA amplification. Full-length single-cell cDNAs are processed for long-read sequencing, while matched aliquots are fragmented for short-read sequencing. Barcodes and transcript sequences are recovered and quantified using the Bagpiper analysis pipeline. (B) Violin plots showing the number of unique molecular identifiers (UMIs) (left) and genes per cell detected per cell (right) and in matched short-read and BenchDrop-seq long-read libraries (n = 2,468 cells). (C) Distribution of equivalence-class lengths derived from Bagpiper quantification, illustrating reduced transcript-assignment ambiguity for single-cell long-read data relative to short-read data. Point size reflects the fraction of reads assigned to each equivalence class. (D) Coverage of the α-globin locus (HBA1), showing pronounced 3’ capture bias in short-read data (top) and uniform, full-length transcript coverage in BenchDrop-seq long-read data (bottom). Orange color denotes reads mapping to the reverse-strand, while Blue denotes forward-strand mapping reads. (E-G) Gene-level pseudobulk expression correlations comparing BenchDrop-seq long-read and bulk RNA-seq (E), short-read and bulk RNA-seq (F), and BenchDrop-seq long-read and short-read RNA-seq (G). Each point represents a gene; color indicates point density. Spearman correlation coefficients are indicated. (H) Subsampling analysis of single-cell long-read data showing the stability of gene-level correlations with bulk RNA-seq across decreasing fractions of the original sequencing depth. Violin plots summarize the distribution of correlations across genes at each subsampling level.

Oxford Nanopore sequencing of BenchDrop-seq libraries produced long-read sequences with a mean length of 660 bp (median, 620 bp; N50, 722 bp; maximum >10 kb), consistent with the expected size distribution of full-length single-cell cDNA (Supplementary Fig. 2A). Long-read libraries showed higher alignment rates to both the human genome (96.0%) and transcriptome (88.6%) than matched short-read libraries (85.7% and 74.8%, respectively; read length 68 bp). Across libraries, we generated 22.7 million long reads and 61.9 million short reads, corresponding to an average of 4,500 and 9,300 unique molecular identifiers (UMIs) per cell across 1400 and 2300 genes, respectively, after filtering and deduplication (Fig. 2B).

Single-cell long-read coverage spanned multiple exons per molecule, substantially reducing transcript-assignment ambiguity relative to short-read data. Nearly one-third of short-read fragments were compatible with more than six transcript isoforms, reflecting limited positional information from 3’-end-biased sequencing. In contrast, BenchDrop-seq long-read alignments typically resolved to fewer compatible transcripts, corresponding to an approximately 60% reduction in isoform-assignment ambiguity quantified by the lengths of equivalence-class (Fig. 2C; Material & Methods).

Using Bagpiper for barcode recovery and isoform-aware gene quantification, we recovered valid cellular barcodes from >90% of single-cell long-read sequences and detected >90% of the isoforms identified in bulk RNA-seq datasets (Material & Methods-Barcode identification). This barcode recovery rate falls within or above the range reported for microfluidics-based single-cell long-read RNA-seq workflows (typically ∼60-75% under comparable filtering criteria^24^), indicating robust barcode assignment despite the elevated per-base error rates associated with long-read sequencing.

As a biological illustration of full-length transcript recovery, we examined the α-globin gene cluster (HBA1, HBA2, HBZ), which is robustly expressed in K562 cells. At this locus, short-read libraries exhibited the expected 3’ capture bias, whereas BenchDrop-seq long-read data spanned the full gene bodies and recovered complete isoforms and splice junctions (Fig. 2D; Supplementary Fig. 2B-C).

Gene-level pseudobulk expression estimates derived from BenchDrop-seq single-cell long-read data were highly concordant with bulk RNA-seq measurements (Spearman’s ρ = 0.86), exceeding correlations observed for matched short-read data (ρ = 0.80; Fig. 2E, F). Concordance between BenchDrop-seq and short-read estimates was also high (ρ = 0.83; Fig. 2G). This agreement persisted when long-read data were subsampled to 25% of the original sequencing depth, indicating stable gene-level quantification at reduced read depth (Fig. 2H). Gene-abundance estimates produced by Bagpiper were also consistent with those obtained using established short-read pipelines, including Salmon-Alevin^25^ and Kallisto-Bustools^26^ (Supplementary Fig. 2D-E).

Together, these analyses establish that BenchDrop-seq supports accurate and reproducible gene-level quantification while reducing transcript-assignment ambiguity relative to short-read single-cell RNA-seq.

### Single-cell gene-level quantification in heterogeneous primary tissue

We next evaluated BenchDrop-seq in human peripheral blood mononuclear cells (PBMCs), a heterogeneous primary tissue comprising multiple immune cell populations. In contrast to the controlled K562 system, this analysis focused on assessing gene-level quantification accuracy and cell-type resolution at single-cell resolution in a complex biological mixture (Fig. 2A-PBMC). Using freshly isolated PBMCs, we captured and processed 19,051 high-quality cells, generating 145.4 million single-cell long-read sequences (N50 = 713 bp; Supplementary Fig. 3A). Of these, 85.1% aligned to the human genome and 64.5% to the transcriptome, yielding a mean of approximately 2,800 deduplicated unique molecular identifiers per cell (Supplementary Fig. 3B).

To assess quantitative accuracy in this heterogeneous setting, we first compared gene-level pseudobulk expression estimates derived from BenchDrop-seq to matched short-read single-cell RNA-seq data and publicly available bulk PBMC RNA-seq datasets. BenchDrop-seq gene-level measurements showed strong concordance with bulk RNA-seq (Spearman’s ρ = 0.85), exceeding correlations observed for matched short-read data (ρ = 0.75; Supplementary Fig. 3C, D). While most genes exhibited broadly consistent expression across platforms, the largest discrepancies reflected systematic biases intrinsic to short-read, end-biased sequencing, whereas BenchDrop-seq measurements showed improved agreement with bulk RNA-seq for these genes.

To determine the sources of these discrepancies, we systematically compared the fold changes in bulk and short reads with those in bulk and long-read-based pseudobulk gene abundance estimates (Fig. 3A; Materials and Methods-Correlative analyses). Genes over-quantified by both single-cell platforms relative to bulk, including RPS4Y1, DDX3Y, and PRKY, were predominantly Y chromosome-encoded and reflected a donor sex mismatch between datasets, serving as an internal positive control. Conversely, histone genes such as HIST1H4B, HIST1H4F, and HIST2H2AB were under-quantified across platforms, consistent with their lack of poly(A) tails and reduced expression in the 3′-biased protocol^27^.

**Figure 3.**
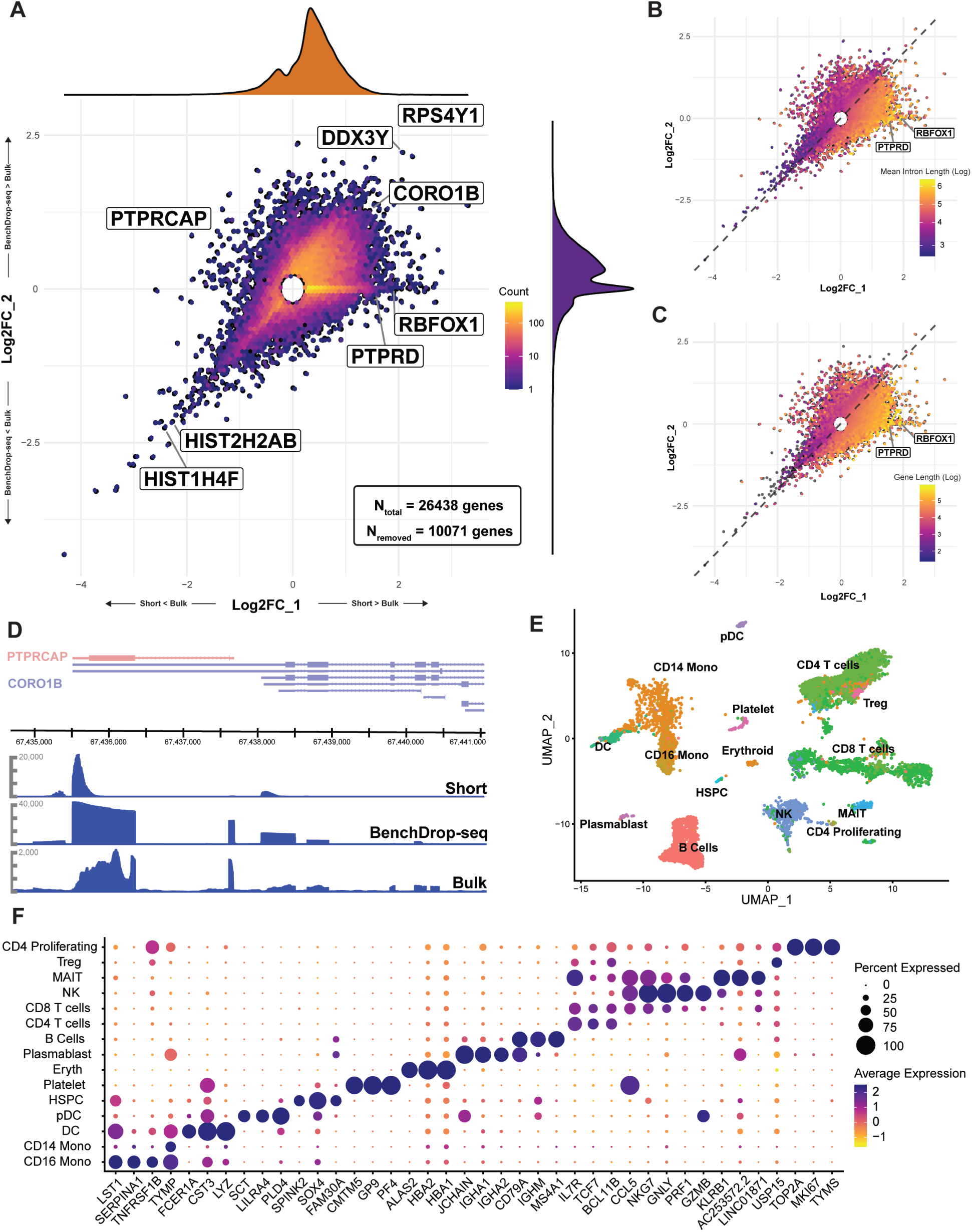
Gene-level quantification accuracy and cell-type resolution in human PBMCs. BenchDrop-seq preserves accurate gene-level quantification in a heterogeneous immune mixture while reducing short-read-specific biases and supporting robust immune cell-type resolution. (A) Scatterplot comparing gene-level log₂ fold changes between short-read single-cell RNA-seq pseudobulk and bulk RNA-seq (x-axis) versus BenchDrop-seq long-read pseudobulk and bulk RNA-seq (y-axis). Each point represents a gene; points are binned (n = 100 genes per bin) and colored by local density. Histograms along each axis indicate the marginal distributions. The central region was removed to enhance visualization of the tails (N_total = 26,438 genes; N_removed = 10,071 genes). Representative genes illustrating platform-specific deviations are annotated. (B) Same comparison as in (A), with bins colored by mean intron length (log₁₀ scale), calculated as the total intronic length divided by the number of introns per gene. Genes with longer introns show increased deviation in short-read-based estimates. (C) Same comparison as in (A), with bins colored by gene length (log₁₀ scale). Increased gene length is associated with systematic over-quantification in short-read data but not in BenchDrop-seq measurements. (D) Genome browser view of the overlapping PTPRCAP and CORO1B loci. Short-read coverage exhibits ambiguity at shared 3’ regions, whereas BenchDrop-seq long-read coverage resolves the loci unambiguously, consistent with bulk RNA-seq profiles. (E) UMAP embedding of BenchDrop-seq gene-level expression profiles from PBMCs, with cells colored by immune cell type transferred from a CITE-seq reference. Major immune populations, including T cell subsets, B cells, natural killer cells, monocytes, dendritic cells, and plasmablasts, are resolved. (F) Dot plot showing expression of canonical marker genes across annotated immune cell types. Dot size indicates the fraction of cells expressing each gene, and color indicates average scaled expression. Marker genes were selected by filtering for avg_log2FC > 0.4 and expressed in at least 25% of the cell type of interest.

Notably, a distinct class of genes was selectively over-quantified in short-read data but showed expression estimates more consistent with bulk RNA-seq when measured using BenchDrop-seq. These genes frequently exhibited overlapping 3’ loci or extended gene lengths, exacerbating ambiguity under short-read coverage constraints. For example, the gene body of PTPRCAP lies entirely within CORO1B, leading to ambiguous short-read assignment at shared 3’ regions, whereas single-cell long-read coverage resolves these loci unambiguously (Fig. 3D). Across genes, short-read over-quantification scaled with increasing gene and intron length, consistent with ambiguity arising from end-biased coverage, whereas this trend was markedly attenuated in BenchDrop-seq data (Fig. 3B-C).

To confirm that BenchDrop-seq supports standard single-cell analyses, we mapped cell-type annotations from a CITE-seq PBMC reference onto BenchDrop-seq gene-level profiles. Dimensionality reduction of BenchDrop-seq expression data resolved all major immune populations, including CD4 and CD8 T cells, B cells, natural killer cells, CD14 and CD16 monocytes, and dendritic cells (Fig. 3E), with appropriate enrichment of canonical marker genes (Fig. 3F).

Together, these results demonstrate that BenchDrop-seq supports accurate gene-level quantification and reliable cell-type resolution in a complex primary immune tissue, while mitigating systematic biases associated with short-read single-cell RNA sequencing.

### Transcript quantification across heterogeneous cell populations

To evaluate transcript-level resolution in a complex biological setting, we applied isoform-aware quantification using Bagpiper to single-cell long-read RNA-seq data generated from human PBMCs. Across 19,051 cells, we identified 2,367 genes exhibiting multiple transcript isoforms with reproducible differential transcript usage across immune cell populations, indicating widespread transcript-level heterogeneity detectable at single-cell resolution.

To assess quantitative concordance at the transcript level, we compared isoform-fraction estimates from BenchDrop-seq with those from bulk long-read RNA-sequencing. Overall agreement was moderate (median Spearman’s ρ = 0.55), consistent with the known challenges of estimating transcript abundances in sparse single-cell datasets^28,29^. This level of concordance reflects a combination of biological variability, reduced per-transcript coverage, and differences in modeling assumptions between bulk and single-cell measurements, rather than a systematic bias in single-cell long-read quantification.

We next compared BenchDrop-seq isoform estimates to those obtained using MAS-ISO-seq^18^, an independent long-read isoform quantification framework applied to PBMC datasets. Across detected transcript isoforms, concordance exhibited a bimodal distribution (Supplementary Fig. 4): approximately half of the isoforms showed stronger agreement between BenchDrop-seq and MAS-ISO-seq than with bulk measurements, whereas the remaining isoforms were more closely correlated with bulk long-read data than with MAS-ISO-seq. This reciprocal pattern illustrates the intrinsic difficulty of transcript-level quantification in single cells and indicates that different long-read analysis frameworks recover overlapping but partially distinct subsets of isoform-level signal, influenced by transcript structure, expression levels, and cell-type specificity. These results highlight the importance of orthogonal comparisons when interpreting transcript-level single-cell measurements.

Despite these quantitative limitations, transcript-level measurements from BenchDrop-seq revealed reproducible, cell-type-specific patterns consistent with established immune cell transcriptional programs. Visualization of transcript-level expression relative to gene-level abundances reveals distinct localization of specific transcript isoforms within defined immune populations. For example, alternative transcript usage of IL7R was preferentially enriched within T cell (CD4^+^CD8^+^) populations, whereas distinct CD8A and NKG7 transcript isoforms showed restricted usage within cytotoxic T cell and natural killer cell populations, respectively (Fig. 4A-C). Similarly, transcript-level heterogeneity at CST3 and SERPINA1 distinguished myeloid populations (Monocytes and Dendritic cells respectively), while CD79A transcript usage was confined to B cell subsets (Fig. 4D-F). Unsupervised clustering of transcript-level counts and Feature plots of individual transcript isoforms further demonstrated that these patterns were not driven solely by gene-level expression, but reflected differential usage of specific transcript structures within cell types (Supplementary Fig. 5A-G).

**Figure 4.**
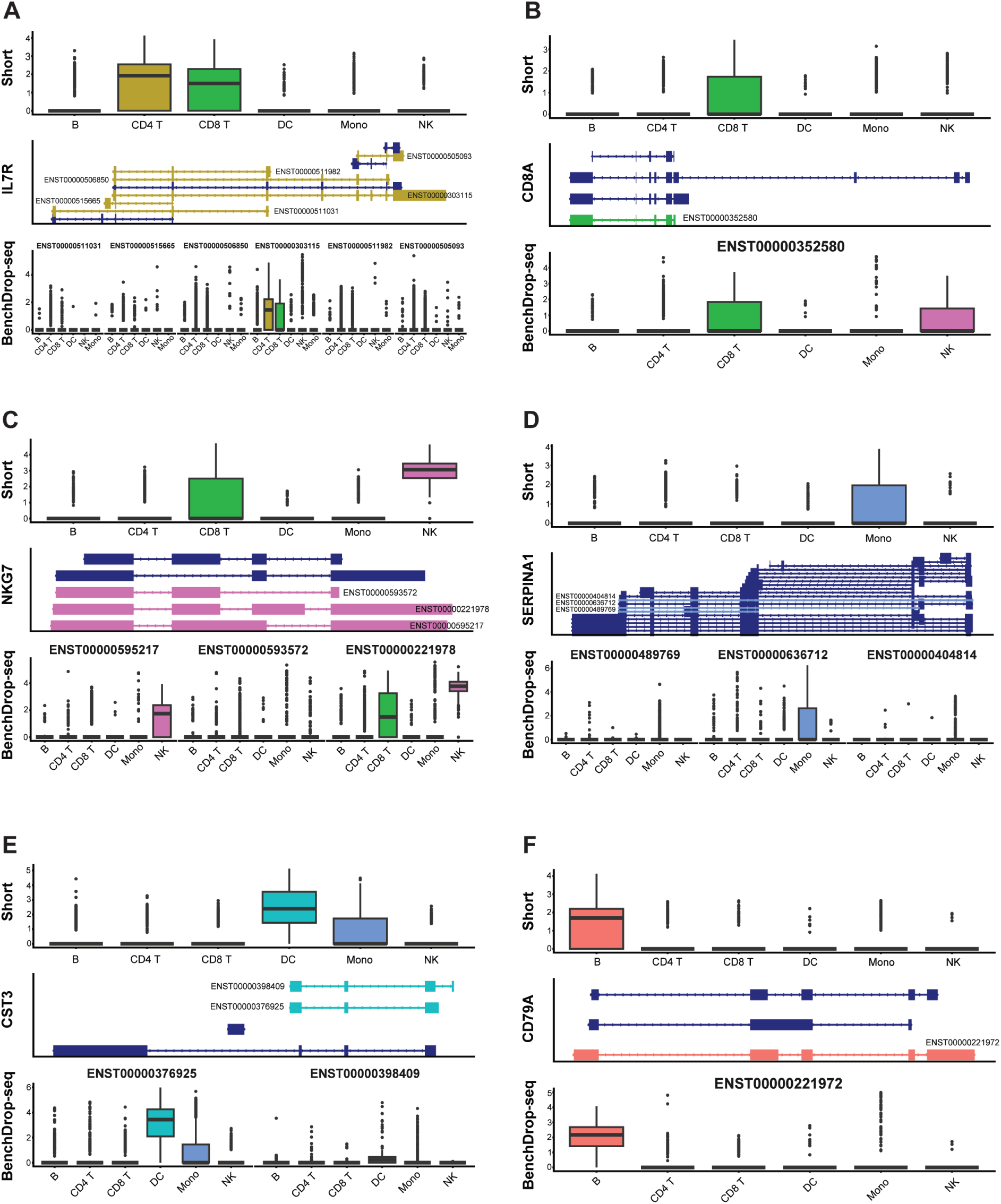
Cell-type-specific transcript usage revealed by single-cell long-read RNA sequencing in PBMCs. BenchDrop-seq resolves reproducible, cell-type-specific transcript usage patterns that are not apparent from short-read, gene-level measurements. Transcript-level expression of (A) IL7R, (B) CD8A, (C) NKG7, (D) SERPINA1, (E) CST3, and (F) CD79A across major PBMC populations. Boxplots compare short-read gene-level estimates (top) with BenchDrop-seq transcript-resolved measurements (bottom), alongside annotated transcript structures. Violin plots are shown only for transcripts that had an average expression above 0.005 across all cells in the BenchDrop-seq library. Across all panels, transcript structures are shown schematically with Ensembl transcript identifiers. Boxplots summarize log-normalized expression across annotated cell types. These examples illustrate how isoform-resolved single-cell measurements capture biologically meaningful transcript usage patterns that complement gene-level analyses and are inaccessible to short-read assays.

Together, these analyses demonstrate that BenchDrop-seq enables detection of biologically meaningful transcript-level heterogeneity in complex primary tissues. While transcript-level quantification remains inherently challenging in single-cell settings, BenchDrop-seq captures transcript usage patterns that complement gene-level analyses and are not readily accessible to short-read assays, delineating both the current capabilities and limitations of transcript-resolved single-cell transcriptomics.

## Discussion

BenchDrop-seq expands the methodological landscape of single-cell transcriptomics by enabling isoform-resolved profiling without reliance on microfluidic instrumentation. Conventional short-read single-cell assays have been instrumental in defining cellular diversity but inherently collapse transcript isoforms into aggregated gene-level measurements, limiting direct interrogation of post-transcriptional regulation. By replacing droplet-based microfluidic partitioning with particle-templated instant partitioning and integrating this workflow with Oxford Nanopore sequencing, BenchDrop-seq enables full-length transcript capture entirely on the benchtop using standard laboratory equipment. Coupled with the open-source Bagpiper analysis pipeline, this framework provides an accessible and reproducible approach to single-cell long-read RNA sequencing.

Benchmarking across a controlled cell line and a heterogeneous primary tissue demonstrates that BenchDrop-seq supports accurate gene-level quantification while substantially reducing transcript-assignment ambiguity relative to short-read approaches. Importantly, this performance is maintained at modest sequencing depth, underscoring the efficiency of single-cell long-read measurements when combined with full-length molecular barcoding. Together, these results indicate that single-cell long-read RNA sequencing can be deployed without prohibitive cost or specialized infrastructure, addressing key practical barriers that have limited broader adoption.

Beyond gene-level accuracy, BenchDrop-seq enables transcript-level analyses that are inaccessible to short-read assays. Isoform-resolved measurements in peripheral blood mononuclear cells reveal reproducible, cell-type-specific transcript usage patterns and refine cellular structure beyond gene-based profiles. These analyses are presented as demonstrations of technical capability rather than comprehensive biological characterization, emphasizing the methodological contribution of the platform.

Several limitations remain. As with other poly(A)-based protocols, BenchDrop-seq is restricted to polyadenylated transcripts and does not capture non-polyadenylated RNA species. Although long-read sequencing has historically been associated with reduced barcode recovery due to elevated error rates, our results indicate that barcode recovery is primarily constrained by library structure and alignment strategy rather than read length. In addition, while long-read sequencing yields fewer reads per flow cell than short-read platforms, comparable gene- and isoform-level quantification can be achieved with substantially fewer reads per cell. Future extensions incorporating targeted enrichment, direct RNA sequencing, or adaptive sampling strategies may further improve sensitivity and transcript coverage.

Together, BenchDrop-seq and Bagpiper establish a practical, microfluidics-free framework for isoform-resolved single-cell transcriptomics. By lowering experimental and computational barriers, this approach enables routine interrogation of transcript isoform diversity in single cells and provides a foundation for broader adoption of long-read technologies in single-cell research.

## Material & Methods

### 1. Experimental Methods

#### 1.1 Cell lines and primary samples

##### K562 cell culture conditions

K562 cells were obtained from the American Type Culture Collection (ATCC). Cells were cultured in L-glutamine pre-supplemented RPMI 1640 medium (Corning,10-040-CM), supplemented with 10% fetal bovine serum (FBS; GeminiBio,100-106), and 1% penicillin-streptomycin (Gibco,15140). Cultures were maintained at 37°C in a humidified incubator with 5% CO₂.

##### K562 sample preparation

K562 cells were harvested and washed twice with 1X DPBS (Corning, 21-031-CM), then pelleted at 400 × g for 10 min at 20°C using a swinging bucket centrifuge. Cell pellets were resuspended in Fluent Cell Suspension Buffer (FB0002440) and analyzed for cell count, clumping, and viability using an AO/PI stain (92% viability) on a LUNA-FL (Logos Biosystems).

##### PBMC sample source & preparation

Healthy human donor blood was obtained from The Wistar Institute’s Phlebotomy Core in compliance with all relevant ethics regulations and IRB-approved protocol (no. 21909324). PBMC isolation from healthy donor blood was performed by layering a 1:1 mixture of heparinized blood and 1X DPBS without calcium or magnesium (Corning, 21-031-CM) over Ficoll-Paque™ PLUS (Cytiva, 17-1440-02), followed by centrifugation at 400 × g for 30 minutes at room temperature with no brake. The PBMC layer was transferred to fresh 50 mL tubes, washed in 1X DPBS containing 2% FBS, and centrifuged at 120 × g for 10 minutes at room temperature with no brake. The cell pellet was then treated with a 5 mL ACK lysing buffer (Quality Biological, 118-156-101) for 5 minutes, followed by two washes in 1X DPBS with 2% FBS and centrifugation at 120 × g for 10 minutes at room temperature with no brake. A final wash was performed in 1X DPBS without FBS, with centrifugation at 400 × g for 10 minutes at room temperature with the brake on. PBMCs were resuspended in Fluent Cell Suspension Buffer (FB0002440), and cell count, clumping, and viability were assessed using AO/PI staining (90% viability) on a LUNA-FL (Logos Biosystems).

#### 1.2 Single-cell capture, cDNA synthesis, and whole transcriptome amplification

Single-cell capture was performed using the Fluent PIPseq platform. 5000, K562 cells, and 40,000 peripheral blood mononuclear cells (PBMCs) were loaded directly into Fluent BioSciences PIP bead reagent mixes of T2 (PIPseq T2 3’ Single Cell RNA Kit v4.0 PLUS) and T20 (PIPseq T20 3’ Single Cell RNA Kit V4.0 PLUS), respectively. 40U Protector RNase Inhibitor (Sigma-Aldrich, RNAINH-RO) was added per reaction. Cell capture, lysis, cDNA synthesis, and whole transcriptome amplification (WTA) were performed according to the manufacturer’s instructions. Briefly, cells were encapsulated in emulsions, lysed, and mRNA was captured on PIP beads before emulsion breaking and reverse transcription. WTA was performed using 12 cycles for T2 (K562) samples and 16 cycles for T20 (PBMC) samples. Amplified cDNA was recovered from PIP beads, SPRI-purified, and quantified using Qubit fluorometer (Thermo Fisher Scientific). cDNA quality and size distribution were assessed using High Sensitivity D5000 Screen Tape on 4200 Tapestation (Agilent Technologies). The synthesized cDNA from K562 cells was used for assay optimization.

#### 1.3 BenchDrop-seq long-read library preparation and Nanopore sequencing

Long-read libraries were prepared for Oxford Nanopore sequencing following the protocol “Ligation Sequencing V14 - Single-Cell Transcriptomics with 3′ cDNA prepared using 10x Genomics on PromethION (SQK-LSK114)”, with modifications to enable amplification of PIP-derived cDNA for Nanopore sequencing. Briefly, 10ng of K562 and 10ng of PBMC cDNA were amplified using the following

Custom biotinylated oligos:

/5Biosg/CAGCACTTGCCTGTCGCTCTCTTTCCCTACACGACGCTCTTCCGATCT and

Partial TSO oligo:

CAGCTTTCTGTTGGTGCTGATATTGCAAGCAGTGGTATCAACGCAGAG

Custom oligos obtained from IDT Technologies are designed to enable efficient, compatible amplification of PIP cDNA. PCR amplification was performed under the following cycling conditions: 94°C for 3 minutes, followed by 4 cycles of 94°C for 30 seconds, 66°C down to 58°C (0.2C/s) for 40 seconds, 58°C for 50 seconds, 65°C for 6 minutes, followed by final extension at 65°C for 10 minutes. Amplified cDNA was purified using 0.8X AMPure XP beads (Beckman Coulter, A63880). Biotinylated fragments were subsequently captured and purified, followed by a second round of PCR using the following custom oligonucleotides:

5’/Phos/CTCTTTCCCTACACGACGCTC and

5’/Phos/AAGCAGTGGTATCAACGCAGAGT for K562 and

5’/Phos/NNNCTCTTTCCCTACACGACGCTC and

5’/Phos/NNNAAGCAGTGGTATCAACGCAGAGT for PBMC cDNAs.

PCR amplification was performed under the following cycling conditions: 94°C for 3 minutes, followed by 4 cycles of 94°C for 15 seconds, 56°C for 15 seconds, and 65°C for 6 minutes, then a final extension at 65°C for 10 minutes. PCR products were purified using 0.8X AMPure XP beads. cDNA concentration was quantified using a Qubit fluorometer (Thermo Fisher Scientific), and fragment size distributions were assessed using the TapeStation system (Agilent Technologies). For library preparation, 200 fmol of K562 cDNA and 200 fmol of PBMC cDNA were end-repaired and adapter-ligated according to the Oxford Nanopore SQK-LSK114 kit protocol. K562 libraries were eluted in 15 µL (volume compatible with MinION) of elution buffer, while PBMC libraries were processed according to the standard protocol. 100fmol of K562 and 100fmol of PBMC libraries were loaded and sequenced on a MinION flow cell (Oxford Nanopore, FLO-MIN114 R10.4.1) on the MinION Mk1D (Oxford Nanopore) and PromethION flow cell (Oxford Nanopore, FLO-PRO114M R10.4.1) on the PromethION 2 Solo (Oxford Nanopore), respectively.

#### 1.4 BenchDrop-seq short-read library preparation and Illumina sequencing

Libraries were prepared for Illumina sequencing and were generated following the guidelines provided with the PIPseq T2 3’ Single-Cell RNA Kit v4.0 PLUS for K562 samples and the PIPseq T20 3’ Single-Cell RNA Kit v4.0 PLUS for PBMC samples. Briefly, 150 ng of amplified K562 cDNA and 100 ng of amplified PBMC cDNA were used as input for PIP-seq short-read library construction. cDNA underwent enzymatic fragmentation, end repair, 3’ A-tailing, and adapter ligation, followed by sample indexing PCR amplification, 7 cycles for K562 libraries and 8 cycles for PBMC libraries. Final libraries were quantified using a Qubit fluorometer (Thermo Fisher Scientific), and fragment size distributions were evaluated using High Sensitivity D5000 ScreenTape on 4200 TapeStation (Agilent Technologies). Indexed libraries were sequenced on an Illumina NextSeq 2000 platform using standard paired-end sequencing chemistry.

#### 1.5 Experimental cost comparison

Across all stages of library preparation, BenchDrop-seq reduced per-cell cost relative to microfluidics-based workflows by minimizing barcoding and capture overhead while maintaining comparable sequencing requirements. For ∼10,000 cells, the total capture-to-sequencing cost per cell was 0.176 USD for BenchDrop-seq, compared with 0.310-0.458 USD for alternative platforms. Importantly, BenchDrop-seq achieved these reductions without increasing protocol duration, requiring approximately 2-3 days from cells to sequencing, comparable to existing approaches (Extended Data Table 1).

To compare the experimental costs and protocol durations across single-cell long-read RNA-sequencing workflows, we estimated per-cell costs using standardized assumptions and publicly available reagent pricing. All cost calculations were performed assuming approximately 10,000 input cells and equivalent sequencing depth across platforms, enabling relative comparison of upstream experimental requirements rather than absolute sequencing yield.

Costs were partitioned into three major stages: (i) cell capture and molecular barcoding through cDNA generation, (ii) long-read library preparation from amplified cDNA, and (iii) sequencing. Reagent costs for cell capture and barcoding were estimated based on list prices for commercial kits or consumables required for each workflow, excluding capital equipment amortization and labor. For BenchDrop-seq, capture and barcoding costs reflect bead-based partitioning reagents and enzymes used in the particle-templated instant partitioning workflow. For microfluidics-based workflows, capture costs do not include the proprietary barcoding machine required for droplet-based encapsulation. For split-pool indexing workflows, capture costs include the cost of repeated indexing reactions and the associated reagents.

Long-read library preparation costs were estimated separately and included biotin pull-down, end repair, adapter ligation, and cleanup steps required for Oxford Nanopore sequencing and the Kinnex protocol. Sequencing costs were assumed to be equivalent across workflows for comparable read depth and were therefore held constant in relative comparisons. All cost estimates reflect reagent-only expenses and are reported on a per-cell basis.

Protocol duration was estimated based on hands-on and incubation times required to progress from input cells to sequencing-ready libraries using published protocols and in-house experience. Reported protocol times reflect typical ranges and do not include optional stopping points. Scalability assessments describe architectural constraints of each workflow, such as dependence on microfluidic devices or split-pool indexing, rather than experimentally validated maximum cell numbers.

A summary of per-cell cost estimates, protocol duration, and qualitative scalability considerations is provided in Extended Data Table 1.

### 2. Computational Methods

#### 2.1 The Bagpiper data processing pipeline

Bagpiper is an open-source computational pipeline designed for processing BenchDrop-seq single-cell long-read RNA sequencing data. The pipeline accepts raw Oxford Nanopore sequencing reads in FASTQ format, along with cellular barcodes and unique molecular identifiers (UMIs). Reference genomes and transcriptomes are provided in FASTA and GTF formats.

The primary outputs of the Bagpiper pipeline include transcript- and gene-level count matrices, isoform equivalence classes, and intermediate alignment files in BAM format. All outputs are generated in standardized formats compatible with downstream single-cell analysis frameworks. An overview of the pipeline architecture is shown in Figure 1 and discussed below.

##### Barcode identification

Cellular barcodes were identified directly from individual long-read sequences using an adaptive Smith-Waterman local alignment strategy. For each read, candidate barcode sequences were aligned against a predefined whitelist using an affine gap penalty model with a scoring scheme empirically optimized for Oxford Nanopore error profiles (match = 2, mismatch = −1, gap open = −1, gap extension = −2). For each read, the highest-scoring barcode alignment was retained. Reads yielding multiple barcode matches exceeding the alignment score threshold were excluded to avoid ambiguous assignments.

BenchDrop-seq libraries follow a structured barcode architecture consisting of four concatenated cellular barcode segments separated by fixed spacer sequences:

Barcode₁ (8 bp) - Spacer₁ (CTCGA) - Barcode₂ (6 bp) - Spacer₂ (CTC) - Barcode₃ (6 bp) - Spacer₃ (CAT) - Barcode₄ (8 bp) - UMI.

The inclusion of invariant spacer motifs provides internal sequence anchors that substantially improve the accuracy of barcode extraction from long-read data with elevated base-calling error rates. During processing, the highest-scoring alignment was used to extract all four barcode segments, which were then matched to corresponding whitelist entries.

Because the proprietary Pipseeker software used in the PIP-seq workflow does not expose barcode whitelist information, barcode whitelists were reconstructed by mapping raw PIP-seq sequencing data to Pipseeker-processed outputs from publicly available datasets. The resulting whitelist barcodes were incorporated into the Bagpiper analysis pipeline and are distributed with the open-source software.

A barcode assignment was considered valid only if all four extracted barcode segments uniquely matched entries in the reconstructed whitelist. Using this approach, valid cellular barcodes were recovered for the majority of long-read sequences, enabling reliable assignment of reads to individual cells for downstream gene- and transcript-level analyses.

##### UMI handling and deduplication

UMIs were extracted from each read based on fixed positional offsets relative to the identified barcode sequence. UMIs were required to match the expected length of 12 nucleotides. UMI deduplication was performed independently for each barcode and transcript using exact UMI matching. To account for Nanopore sequencing errors, UMIs differing by a Hamming distance of one were collapsed. UMI collision rates were assumed to be negligible given the UMI complexity and observed molecule counts per cell. No cross-gene or cross-transcript UMI collapsing was performed.

##### Alignment of long reads

Long-read sequences were aligned to the reference transcriptome using Minimap2 v2.29. Alignments were performed using the -ax map-ont preset with the following parameters: -for-only, -N200, and -p 0.9. The human reference genome GRCh38 and corresponding GENCODE v32 transcript annotation were used for all analyses unless otherwise specified. Secondary and supplementary alignments were retained for downstream isoform-aware quantification.

##### Isoform-aware quantification

Isoform abundance estimation in Bagpiper builds on a generative RNA-seq framework originally introduced by Li et al. (2010) and subsequently adapted for bulk long-read transcript quantification by methods such as Oarfish. Bagpiper extends this class of models to single-cell long-read RNA-seq data using a simplified and computationally efficient formulation tailored to the characteristics of full-length single-cell reads.

Two modifications are central to this extension. First, unlike short-read single-cell data, long-read sequencing captures transcript sequences from both forward and reverse strands, introducing ambiguity when overlapping genes are encoded on opposite strands. Bagpiper explicitly accounts for this by performing strand-aware transcript quantification, ensuring that reads are only considered compatible with transcripts on the correct genomic strand (Supplementary Fig. 1B).

Second, Bagpiper models sequencing reads as arising from a mixture of transcript isoforms and infers relative isoform abundances through iterative probabilistic assignment of reads to compatible transcripts. Rather than relying directly on raw alignment scores, Bagpiper incorporates transcript length information within each equivalence class to modulate read-transcript compatibility. Specifically, the conditional contribution of a read to a candidate transcript is weighted by the ratio of the shortest compatible transcript length to the length of the transcript under consideration, providing a simple and robust proxy for alignment likelihood in the presence of long-read sequencing error.

Isoform abundances are estimated using an expectation-maximization (EM) algorithm. During each iteration, reads are assigned to compatible transcripts in proportion to their current abundance estimates and adjusted compatibility weights, and transcript abundances are updated accordingly. Iterations proceed until convergence, defined as a maximum relative change in transcript abundance below 0.01 across successive iterations, or until a maximum of 100 iterations is reached, consistent with prior single-cell RNA-seq quantification frameworks such as alevin.

##### Gene-level quantification

Gene-level abundance estimates were obtained by summing transcript-level abundance estimates across all transcripts assigned to the same gene. Genes were retained for downstream analyses if they were detected in at least 10 cells and had a total UMI count of at least 100 across the dataset. No additional normalization was applied prior to pseudobulk analyses.

#### 2.2 Cell-type annotation

For all sequencing data, the generated count matrices were processed using the Seurat^30^ v5.3 package in R v4.5. Short-read gene count matrices generated with PIPseeker^15^ v3.3.0 were selected corresponding to the sensitivity 3 output of the program. Long-read gene and isoform count matrices generated with Bagpiper were filtered to match the cellular barcodes in the short-read data. For both the BenchDrop-seq short-read and long-read gene matrices, the data were normalized NormalizeData(), the top 2,000 variable features were identified FindVariableFeatures(), the data were scaled ScaleData(), and both PCA (RunPCA()) and UMAP (RunUMAP()) analyses were performed. For the long-read isoform matrix, the same workflow was applied, except that the top 20,000 variable features were identified and used for downstream analysis. For both the short-read and long-read PBMC data, Azimuth^30^ was used to assign cell type labels using the Azimuth Human PBMC Reference 2.10.

#### 2.3 Correlational Analyses

All correlation analyses were performed using pseudobulk expression profiles. For comparisons between single-cell datasets (e.g., BenchDrop-seq, Pipseeker^15^, Salmon-Alevin^31^, Kallisto-Bustools^26^, Mas-ISO-seq^18^), pseudobulk counts were generated by summing normalized gene counts across all cells within each dataset. The union of genes detected across the two datasets was used, and genes present in only one dataset were assigned a count of 0 in the other.

For comparisons between single-cell and bulk RNA-seq datasets, pseudobulk profiles were obtained by summing normalized gene counts across all cells in the single-cell dataset and subsequently rescaling the resulting counts to counts per million (CPM). As above, genes absent from the union dataset were assigned a value of zero.

For any log-fold-change correlation analyses, values between pseudobulk profiles were computed after adding a pseudocount of 1 to all gene or transcript counts to avoid division by zero.

#### 2.4 Computational Runtime Analyses

All computational analyses were performed on a Linux workstation equipped with Intel(R) Xeon(R) Platinum 8462Y+, and running Ubuntu 22.04.5 LTS. No specialized hardware or cloud-based acceleration was required.

To assess computational performance, we recorded wall-clock runtime for each major Bagpiper module, including barcode identification, UMI handling and deduplication, long-read alignment, and isoform-aware quantification. Runtime measurements were obtained using standard UNIX timing utilities and averaged across multiple runs where applicable.

Barcode identification and error correction scaled linearly with the number of reads and required 12 minutes for the full K562 dataset and 70 minutes for the PBMC dataset.

Long-read alignment with Minimap2 v2.29 accounted for the dominant computational cost, taking 40 minutes (>50% of the overall time) for PBMC-scale datasets. Isoform-aware quantification using the expectation-maximization framework took 11 minutes and used less than 2 GB of memory.

For the K562 datasets comprising approximately 150 million reads, the total processing time was ∼60 minutes, with read alignment using minimap2 accounting for the majority of the runtime, followed by equivalence-class construction and quantification. Barcode recovery and UMI processing contributed a minor fraction of total compute time (Supplementary Fig. 1C).

Runtime scaled approximately linearly with the number of input reads, indicating that Bagpiper does not introduce additional computational bottlenecks beyond standard long-read alignment. These results demonstrate that isoform-aware single-cell quantification using Bagpiper is computationally tractable and compatible with routine use on commodity hardware.

For comparison, short-read single-cell RNA-seq data generated from matched cDNA pools were processed using standard pipelines (Salmon-Alevin v1.10.0 and Kallisto-Bustools v0.51.1) with recommended parameters. Total runtime for short-read pipelines was measured using the same hardware and timing framework. While short-read alignment and quantification were faster per read, overall runtime scaled with the substantially higher read counts required for comparable gene-level resolution. When normalized per cell and per informative molecule, Bagpiper exhibited comparable computational efficiency while enabling transcript-level resolution not accessible to short-read pipelines.

Together, these analyses demonstrate that Bagpiper supports practical single-cell long-read RNA-seq analysis on standard laboratory compute infrastructure, without requiring specialized hardware or prohibitive runtime.

## 3. Software and Data Availability

The Bagpiper data processing pipeline is implemented in Rust and is available as open-source software at https://github.com/avisrilab/bagpiper. The version used for all analyses in this study is v0.1.0, corresponding to commit c196b8b. All analyses reported in this manuscript were performed using this version unless otherwise specified and can be replicated using scripts at https://github.com/JamieBregman/BenchDrop-seq_Analysis.

Bulk RNA-seq datasets used for benchmarking and validation were obtained from the ENCODE Project Consortium (SRX2370564). Gene-level comparisons were performed using consistent gene identifiers and annotation versions across datasets. All datasets were harmonized to the same reference genome, GRCh38, and transcript annotation, GENCODE v32, prior to analysis.

Detailed instructions for reproducing all analyses, including software dependencies and execution commands, are provided in the Bagpiper GitHub repository.

## 4. Author Contribution

R.R. and A.S. conceived the study. J.B. performed computational work supervised by A.S.; C.N. performed experimental work supervised by A.S. R.R drafted figure 1 and supplementary figure 1. R.R. & A.S. wrote the initial draft of the manuscript while all authors participated in the interpretation and revision of the manuscript.

## 5. Acknowledgment

We thank Noam Auslander, Bin Tian, and Simon Chu for their insightful discussions and feedback, which contributed to the development of this work. We also thank the Wistar Cancer Center Genomics Shared Resource (P30-CA010815) for providing outstanding technical support. The graphical abstract was created in BioRender.

