## Supplementary material for "BenchDrop-seq: a microfluidics-free platform for benchtop single-cell long-read RNA sequencing": Supp


| All Numbers for ~10k cells | BenchDrop-seq | 10x + Nanopore | 10x + Kinnex (PacBio) | Parse Bio + Kinnex (PacBio) |
|---|---|---|---|---|
| Capture and Barcoding to cDNA | $X | 3x $X | 3x $X | 5x $X |
| Biotin pull-down to Long-Read Library | $Y | $Y | 3x $Y | 3x $Y |
| Sequencing | $Z | $Z | $Z | $Z |
| Protocol time from cells to loading onto sequencer | ~2-3days | ~2-3days | ~3-4days | ~3-4days |
| Cost per cell (10,000 cells) Capture-Sequencing | 0.176 | 0.310 | 0.3512 | 0.458 |
| Special Considerations: Fixation | Live/Fixed cells | Live/Fixed cells | Fixed cells | Fixed cells |
| Special Considerations: Barcoding | No microfludics / Vortexing | Microfluidics | Microfluidics | Split-pooling |
| Special Considerations: Architecturally scalable beyond 10k | Scalable to 1M | Fixed at ~10k-15k | Scalable to >1M | Scalable to >1M |

**Supplementary Table 1 |** Relative cost comparison of long-read single-cell platforms

Comparison of per-cell cost, protocol duration, and scalability for representative long-read single-cell RNA-sequencing workflows, assuming approximately 10,000 input cells. Costs are estimated relative to BenchDrop-seq for major experimental stages, including cell capture and barcoding, long-read library preparation, and sequencing, using current reagent pricing and publicly available protocols. Protocol duration reflects hands-on time from cells to sequencing-ready libraries. Scalability and special considerations summarize architectural constraints of each approach rather than experimentally validated limits. Sequencing costs were approximately equivalent across platforms at comparable read depths.

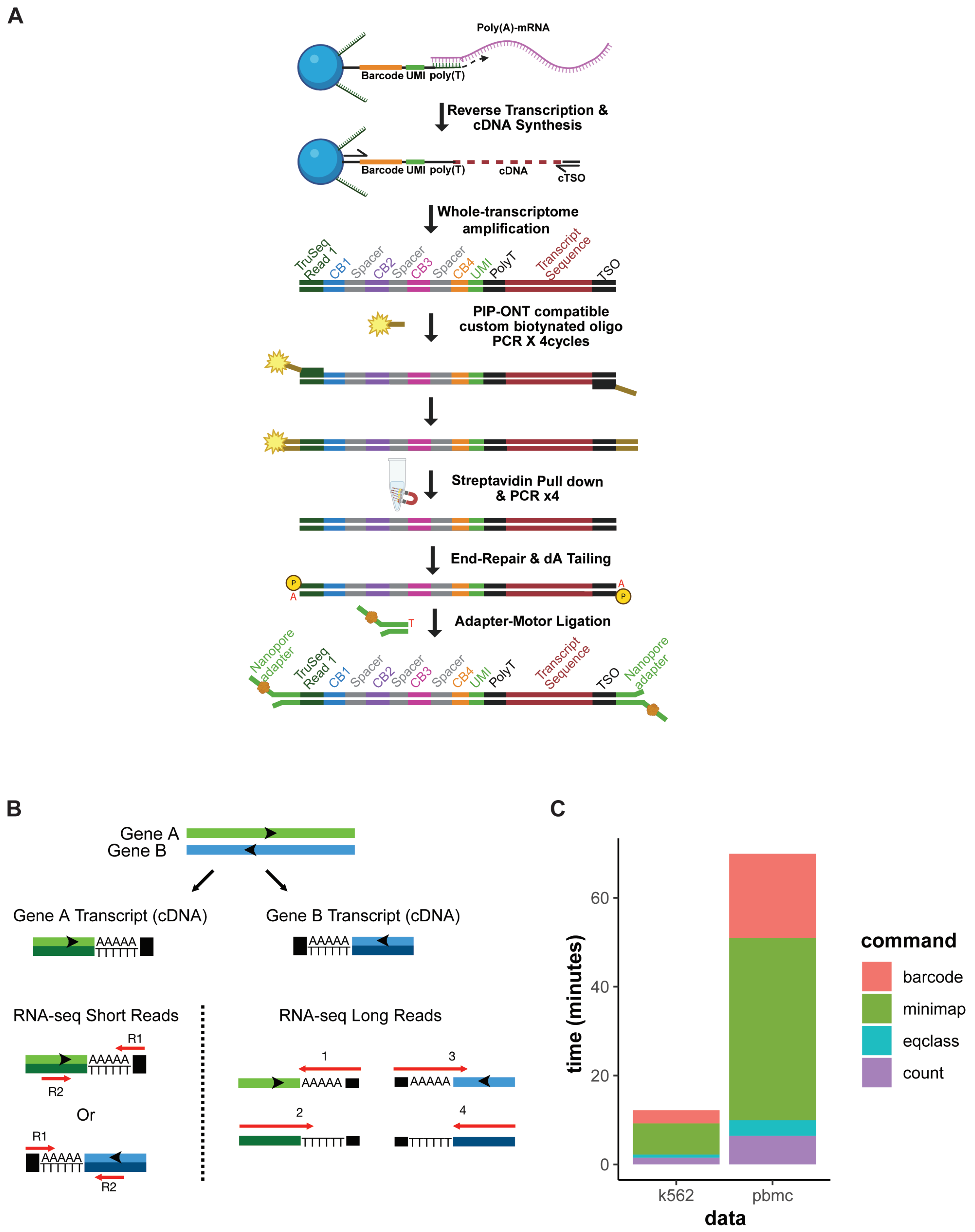


**Supplementary Figure 1** | Detailed characterization of the BenchDrop-seq library structure and computational runtime.

(A) Detailed schematic of the BenchDrop-seq cDNA and library preparation workflow adapted for Oxford Nanopore sequencing. Polyadenylated mRNA molecules are captured on barcoded beads, reverse-transcribed, and then subjected to whole-transcriptome amplification. A custom biotinylated PCR strategy enables selective enrichment of full-length cDNA molecules compatible with Nanopore adapter ligation, preserving the complete barcode-UMI-transcript structure in the final sequencing library.

(B) Conceptual comparison of strand orientation in short-read and long-read RNA sequencing. Short-read libraries preserve strand information but sample only a limited portion of each transcript, leading to ambiguous assignment for overlapping genes on opposite strands when reads originate from shared genomic regions. In contrast, long-read sequencing captures full-length transcript molecules and directly records their strand of origin. This increased structural complexity necessitates explicit strand-aware computational modeling to correctly resolve exon-intron architecture and transcript orientation in long-read data.

(c) Computational runtime of the Bagpiper analysis pipeline for K562 and PBMC datasets, broken down by major processing steps, including barcode identification, read alignment (minimap2), equivalence-class construction, and transcript quantification. Total runtime scales with dataset size and is dominated by long-read alignment, while barcode recovery and quantification contribute a minor fraction of overall compute time.

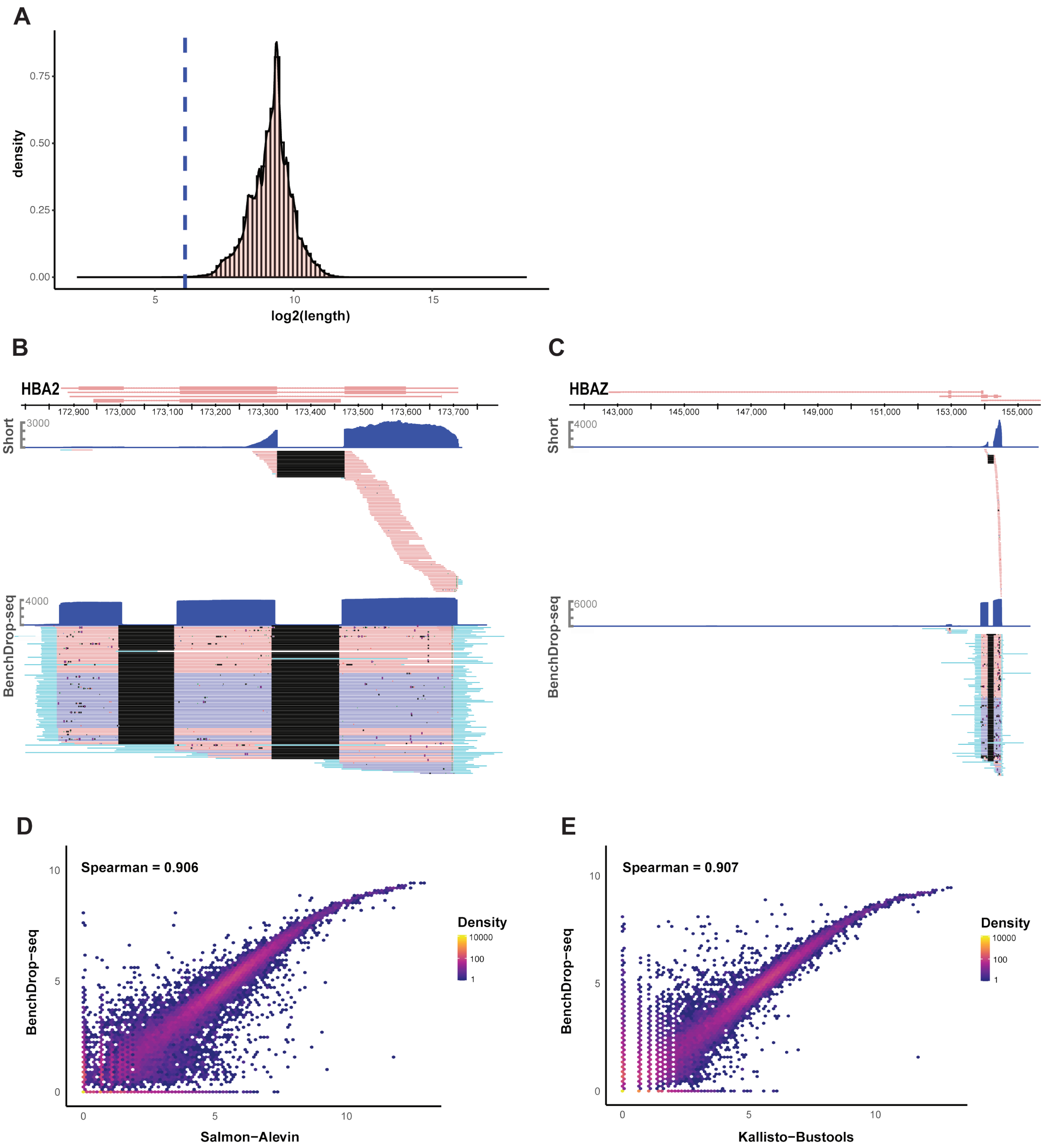


**Supplementary Figure 2 |** Additional validation of long-read coverage, isoform recovery, and gene-level concordance in K562 cells.

(A) Distribution of $log_2$-transformed read lengths (in base-pairs) for BenchDrop-seq single-cell long-read data from K562 cells. The dashed vertical line indicates the minimum read-length threshold for short-read data, illustrating that most long-reads are substantially longer.

(B) Coverage of the HBA2 locus comparing short-read and BenchDrop-seq long-read data. Short-read data show pronounced 3′-end bias, whereas BenchDrop-seq long reads span the full gene body and resolve transcript structure across multiple exons.

(C) Coverage of the HBAZ locus illustrating recovery of structurally distinct transcripts with BenchDrop-seq long-read sequencing relative to short-read data.

(D) Gene-level abundance correlation between BenchDrop-seq long-read data quantified with Bagpiper and short-read data quantified using Salmon-Alevin (Spearman's $\rho = 0.906$). Each point represents a gene; color denotes local point density.

(E) Gene-level abundance correlation between BenchDrop-seq long-read data and short-read data quantified using Kallisto-Bustools (Spearman's $\rho = 0.907$), demonstrating consistency across alternative short-read quantification frameworks.

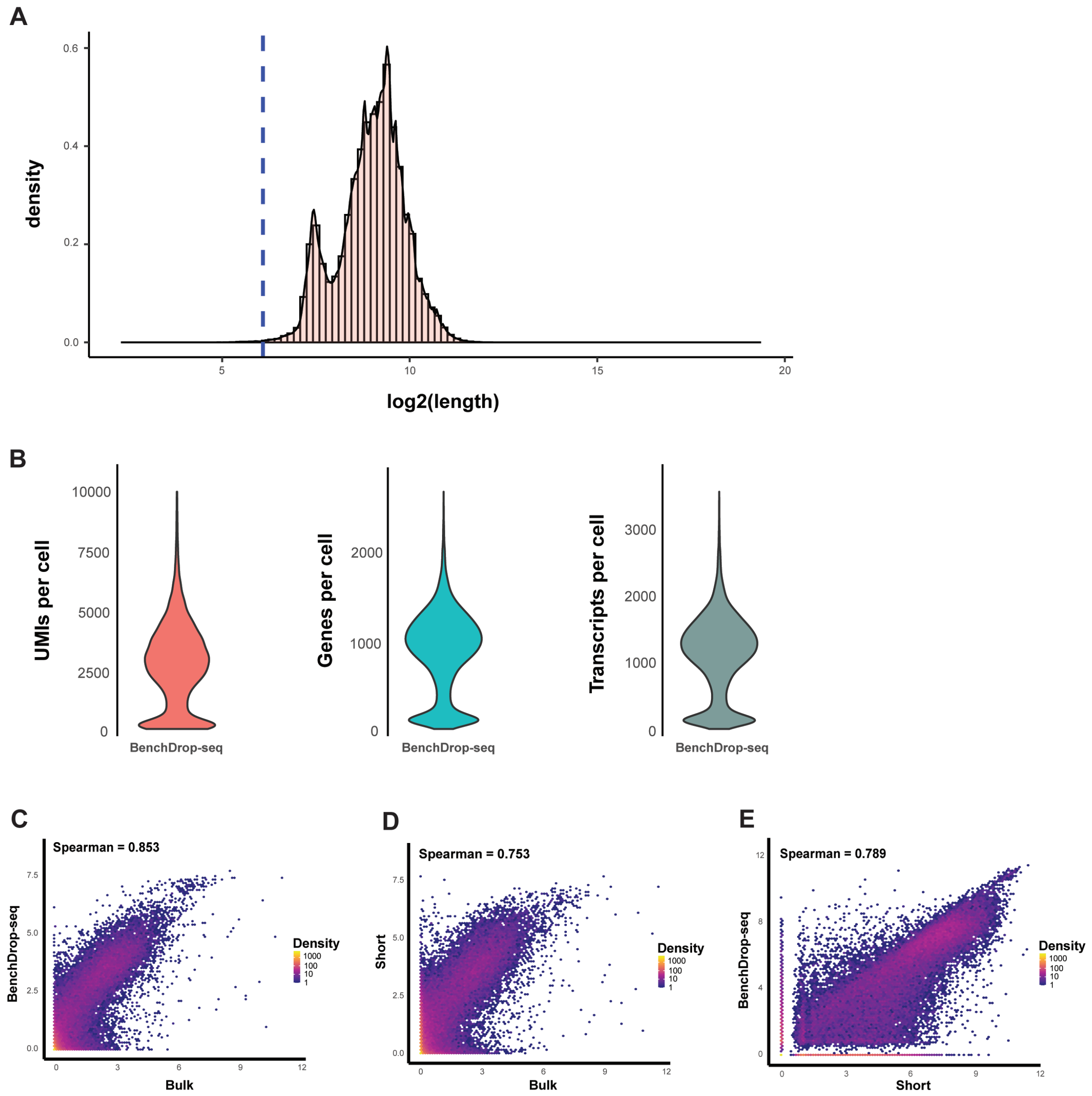


**Supplementary Figure 3 |** Data quality, library complexity, and gene-level concordance in PBMC BenchDrop-seq datasets.

(A) Distribution of single-cell long-read lengths for PBMC BenchDrop-seq libraries, shown as a density histogram of log2-transformed read lengths (in base-pairs). The dashed vertical line indicates the minimum read-length for short-read RNA-seq data.

(B) Violin plots summarizing per-cell library complexity in PBMC BenchDrop-seq data, including the number of unique molecular identifiers (UMIs), genes detected, and transcripts detected per cell after filtering.

(C) Gene-level pseudobulk expression correlation between BenchDrop-seq single-cell long-read data and bulk RNA-seq (Spearman's $\rho = 0.853$), showing improved concordance relative to short-read data.

(D) Gene-level pseudobulk expression correlation between short-read single-cell RNA-seq and bulk RNA-seq for PBMCs (Spearman’s ρ = 0.753).

(E) Correlation between BenchDrop-seq single-cell long-read and matched short-read pseudobulk gene expression estimates (Spearman’s ρ = 0.789).

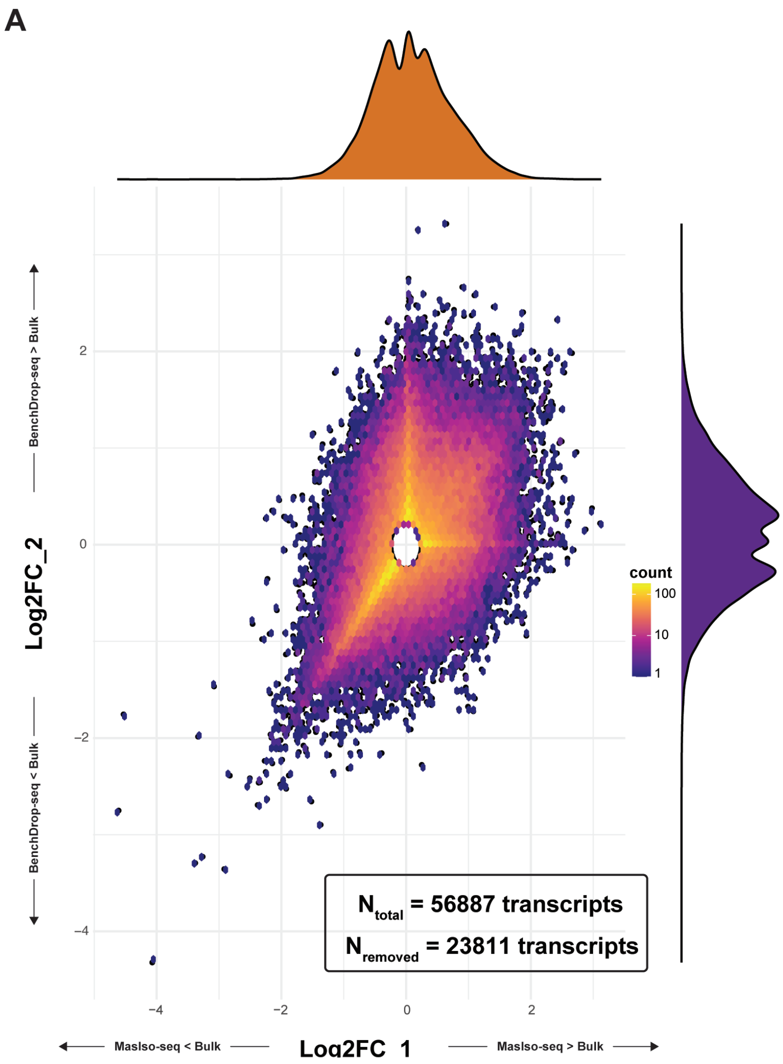


**Supplementary Figure 4 |** Transcript-level concordance across long-read quantification frameworks

(A) Density scatter plot comparing transcript-level log2 fold changes estimated from BenchDrop-seq pseudobulk data relative to bulk long-read RNA sequencing (y-axis) against corresponding estimates from MAS-ISO relative to bulk (x-axis). Each point represents a transcript isoform, with color indicating local point density. Marginal density plots summarize the distribution of log2 fold changes along each axis. Transcripts near the origin, where fold-change estimates are close to zero in both comparisons, were excluded from correlation analyses to reduce the influence of low-information measurements (inset). The remaining transcripts illustrate a broad but structured relationship between isoform-level estimates obtained using different long-read single-cell quantification frameworks, reflecting both shared signal and method-specific variability in sparse single-cell data.

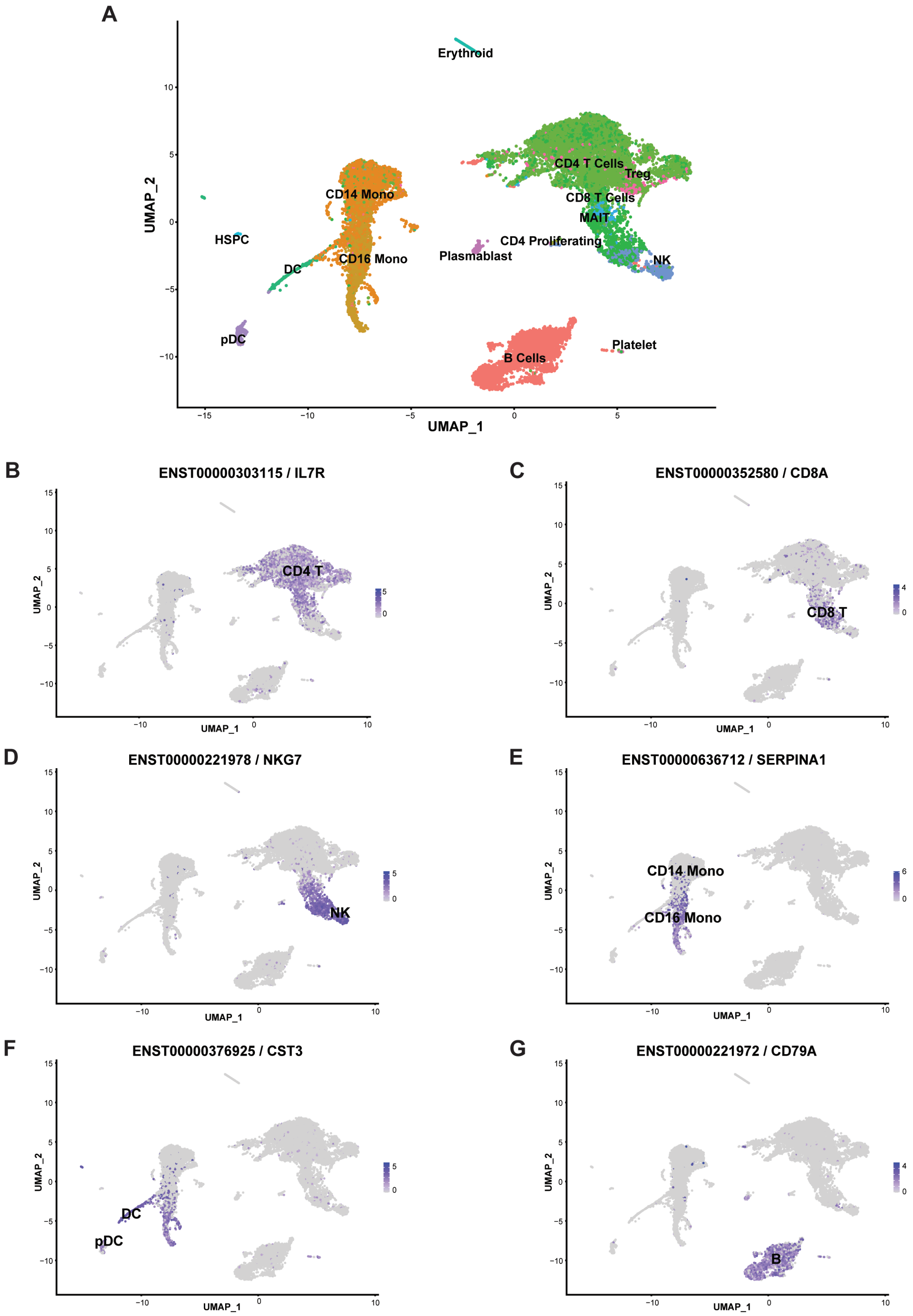

A
Erythroid
CD4 T Cells
Treg
CD8 T Cells
MAIT
CD14 Mono
CD16 Mono
HSPC
DC
pDC
CD4 Proliferating
Plasmablast
NK
B Cells
Platelet
UMAP_1
UMAP_2
B
ENST00000303115 / IL7R
CD4 T
C
ENST00000352580 / CD8A
CD8 T
D
ENST00000221978 / NKG7
NK
E
ENST00000636712 / SERPINA1
CD14 Mono
CD16 Mono
F
ENST00000376925 / CST3
DC
pDC
G
ENST00000221972 / CD79A
B

**Supplementary Figure 5 |** Cell-type-specific transcript usage resolved by single-cell long-read RNA sequencing in PBMCs.

(A) Unsupervised UMAP embedding of transcript-level expression profiles from 19,051 PBMCs profiled by BenchDrop-seq, with cells colored by immune cell type labels transferred from a CITE-seq reference. Major immune populations—including CD4$^{+}$ and CD8$^{+}$ T cells, regulatory T cells, B cells, natural killer (NK) cells, CD14$^{+}$ and CD16$^{+}$ monocytes, dendritic cells (DC and pDC), plasmablasts, platelets, erythroid cells, and hematopoietic stem and progenitor cells (HSPCs)—are clearly resolved.

(B-G) Feature plots showing transcript-level expression of representative immune-associated transcripts projected onto the UMAP embedding. Each panel displays expression of a specific transcript isoform (Ensembl transcript ID shown) quantified from single-cell long-read data.

(B) IL7R transcript ENST00000303115 shows preferential usage within CD4$^{+}$ T cell populations.

(C) CD8A transcript ENST00000352580 is selectively enriched in CD8$^{+}$ T cells.

(D) NKG7 transcript ENST00000221978 is restricted to cytotoxic lymphocytes, including NK cells.

(E) SERPINA1 transcript ENST00000636712 highlights transcript-level heterogeneity within monocyte populations.

(F) CST3 transcript ENST00000376925 is enriched in dendritic cell populations, including pDCs.

(G) CD79A transcript ENST00000221972 shows B-cell-specific transcript usage.

Across all panels, color intensity represents log-normalized transcript abundance, with non-expressing cells shown in grey.
